# Population genetics as a tool to elucidate pathogen reservoirs: Lessons from *Pseudogymnoascus destructans*, the causative agent of white-nose disease in bats

**DOI:** 10.1101/2021.02.04.429503

**Authors:** Nicola M. Fischer, Andrea Altewischer, Surendra Ranpal, Serena Dool, Gerald Kerth, Sébastien J. Puechmaille

## Abstract

Emerging infectious diseases pose a major threat to human, animal, and plant health. The risk of species-extinctions increases when pathogens can survive in the absence of the host, for example in environmental reservoirs. However, identifying such reservoirs and modes of infection is often highly challenging. In this study, we investigated the presence and nature of an environmental reservoir for the ascomycete fungus *Pseudogymnoascus destructans*, the causative agent of white-nose disease. We also characterised the modes and timing of transmission of the pathogen; key elements to better understand the disease dynamics. Using 18 microsatellite markers, we determined the genotypic and genic (based on allele frequencies) differentiation between 1,497 *P. destructans* isolates collected from nine closely situated hibernacula in North-Eastern Germany. One hibernaculum was the focus of intensive sampling in which both the bats and walls of the site were sampled at regular intervals over five consecutive winter seasons (1,062 isolates). We found significant genic differentiation between sites and few multi-locus genotypes shared across hibernacula (genotypic differentiation). This demonstrates that each hibernaculum has an essentially unique population of the fungus. This would be expected if bats purge viable *P. destructans* over the summer, preventing the mixing and exchange of the pathogen in maternity colonies, where bats from all of the studied hibernacula meet. Results from the intensively sampled site show higher measures of genotypic richness on walls compared to bats, the absence of genic differentiation between bats and walls, and stable relative abundance of multi-locus genotypes over multiple winter seasons. This clearly implicates hibernacula walls as the main environmental reservoir of the pathogen, from which bats become re-infected annually.

## Introduction

Emerging infectious diseases pose a major threat to the health of humans, animals and plants, and consequently, to global biodiversity (Daszak et al., 2000; Schmeller et al., 2020). Host-pathogen relationships are ubiquitous in nature. The main reason why these relationships only rarely lead to species extinctions is due to density-dependent transmission, in which transmission decreases when population sizes are low, allowing populations to recover (McCallum, 2012). However, if pathogens can survive in the absence of a host in environmental reservoirs, the risk they pose to biodiversity is markedly increased. This ability is often found in fungal pathogens, including *Batrachochytrium dendrobatidis* (causing amphibian chytridiomycosis; Johnson & Speare, 2003), the *Fusarium solani* species complex (causing disease in a range of hosts, e.g. plants, humans, sea turtles; Zhang et al., 2006) and *Puccinia graminis* (causing wheat stem rust e.g. Rowell & Romig, 1966).

A pathogens’ ability to spread between host populations and survive on alternative hosts or in environmental reservoirs is a key determinant in disease management (De Castro & Bolker, 2004). Unfortunately, it is precisely this information that is often lacking when dealing with newly emerging infectious diseases. This scenario was encountered during the sudden appearance of the ascomycete fungus *Pseudogymnoascus destructans* in North America in 2006, causing the generally lethal White-Nose disease in hibernating bats (Blehert et al., 2009). Although the fungus was first described in North America, it was subsequently found to be present across Europe and Asia (Frick et al., 2016). Population genetic analyses and a lack of mass mortality in European bats indicate that the fungus was introduced from Europe to North America (Drees et al., 2017a; Fritze & Puechmaille, 2018; Leopardi et al., 2015; Puechmaille et al., 2011). In its introduced range, *P. destructans* became an invasive pathogen killing more than 5 million bats within the first 4 years of discovery (www.whitenosesyndrome.org). Due to its threat to bat populations and the ecosystem services they provide (Kunz et al., 2011), there is an urgent need to limit the impact of the disease on bat species. This task is hampered by lack of knowledge about the mode of transmission and potential environmental reservoirs of the fungus (see e.g. Szymanski et al., 2009). Such information is critical for designing adequate management strategies to control the disease (Meyer et al., 2016).

Understanding the life cycle of newly described pathogens emerging in wildlife is a challenging task that takes time. In the case of *P. destructans*, several studies have identified strong links between the host (bats) and the pathogens’ life cycle (Fischer et al., 2020; Langwig et al., 2015a; Puechmaille et al., 2011). Therefore, it becomes important to integrate information about the host behaviour and its life cycle to better understand and characterise the pathogen’s life cycle, including the mode of transmission. Given the strong seasonality of temperate regions, the yearly life cycle of bats living there can be divided into two main seasons (summer and winter), separated by transition periods (spring and autumn). Many temperate insectivorous bats spend the coldest months hibernating in underground sites (i.e., hibernacula). During this period, bats exhibit a reduced body temperature making them susceptible to infection with the cold-loving fungus *P. destructans* (Verant et al., 2012). In spring, bats leave their hibernacula to form summer colonies. The hibernation site is mostly unoccupied until bats return in autumn. During summer, temperate bats are mostly active and females gather in maternity colonies to give birth and raise their pups, while males roost alone or in small groups (e.g. Kunz & Fenton, 2005). Given that *P. destructans* is unable to grow above 20°C and even spores do not survive long periods at elevated temperatures (viability up to 15 days at 37°C), it is most likely that bats clear the infection over the summer period (Kunz & Fenton, 2005; Verant et al., 2012, Campbell et al., 2020). To continue its lifecycle, the fungus must re-infect bats just before or at the onset of the hibernation season (Figure 1). Following the summer breeding period, individuals from several species may meet in pre-hibernation swarming sites (typically also used as hibernacula) where mating occurs. During swarming bats land and crawl on the walls and this behaviour is hypothesised to result in bats becoming infected from the environment (Puechmaille et al., 2011). The fact that *P. destructans* spores persist on hibernacula walls (Vanderwolf et al., 2016) and that the seasonal pattern of *P. destructans* germination is synchronised with the presence of bats in hibernacula (Fischer et al., 2020) further support this inferred mode (wall to bat) and timing of infection (swarming behaviour at swarming/hibernation sites).

**Figure 1.**
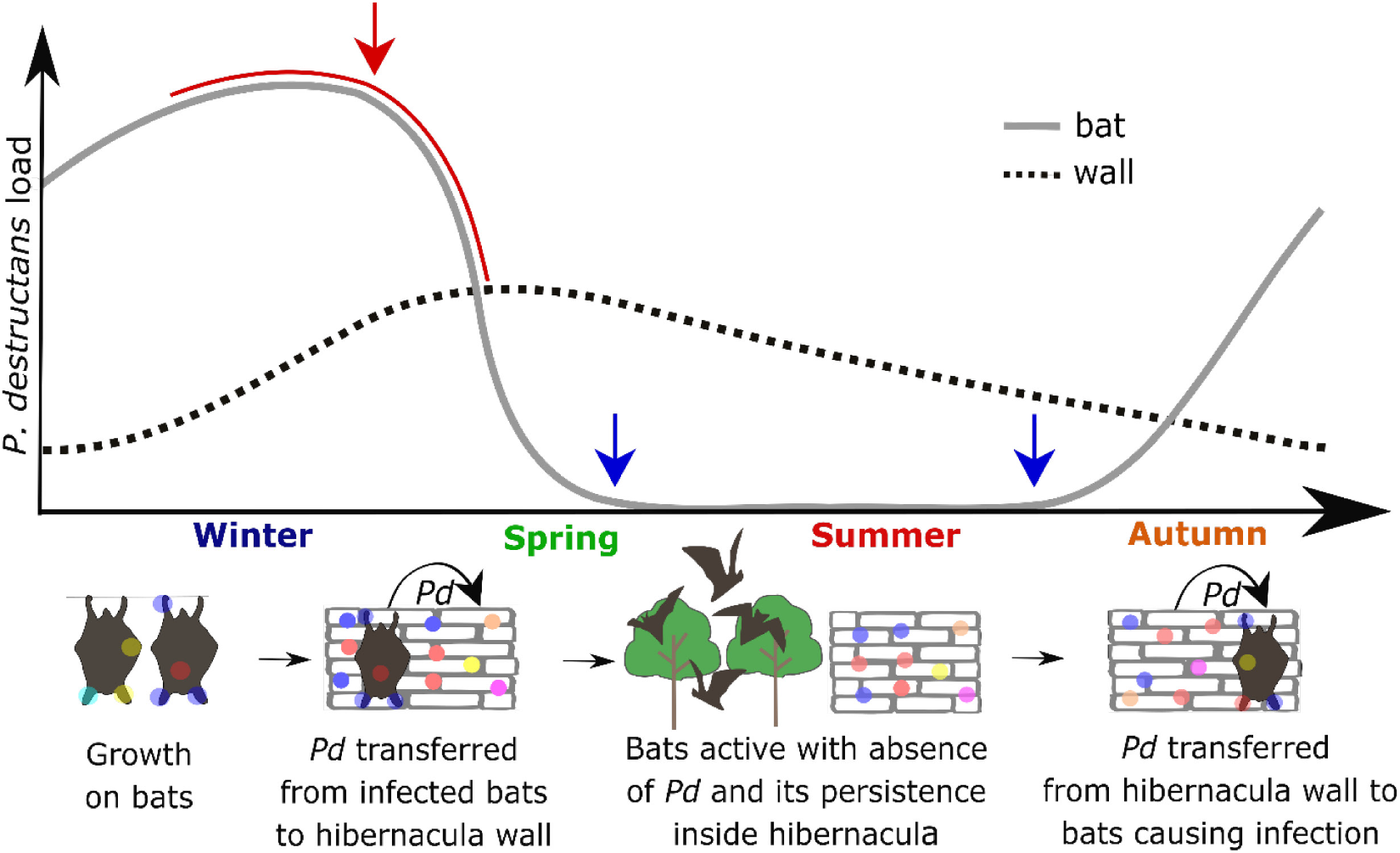
Schematic representation of the life cycle of *P. destructans* (*Pd*) on bats (solid grey line) and in an environmental reservoir (hibernaculum; dotted line) throughout a year as predicted from existing literature. Timing of sampling is indicated by a red line and arrow (bat sampling) and blue arrows (wall sampling). Seasonality of the bat and fungal life cycles are illustrated; coloured dots represent different fungal multi-locus genotypes.

In this study, we analysed populations of *P. destructans* on bats and on their surrounding environment to test the hypothesis that bats’ yearly re-infection originates from an environmental reservoir, namely the walls of their hibernacula. These data were also further analysed to better characterise the transmission pathway. Our work takes advantage of the clonal mode of reproduction of *P. destructans*, enabling us to follow the pathogen through time (summer/winter across several years) and space (environment/bat), which allows the elucidation of transmission routes. Specifically, the pathogen population of a host will be very similar to the pathogen population of the reservoir it became infected from, and more dissimilar to other pathogen populations from which it is not connected via transmission routes. While these comparisons are classically inferred from population differentiation (using allele frequencies), a finer scale approach was additionally used in this study distinguishing fungal individuals (i.e., multi-locus genotypes – MLGs). As fungal individuals cannot be distinguished phenotypically, we used genetic tools to determine the genetic fingerprint of fungal individuals (classically referred to as genets; Burnett, 2003). To accomplish this, we cultured swab samples collected from bats and the hibernaculum walls to obtain fungal single-spore isolates (SSI) which were then genotyped at 18 microsatellite loci used to discriminate MLGs (i.e., individuals).

To address the overall question concerning the presence and nature of the environmental reservoir, we tested successive predictions (see Figure 2). First, if bats clear infection during the summer and are only re-infected in the hibernacula in autumn, bats from different hibernacula would not be able to exchange viable *P. destructans* spores in the summer maternity colonies where they meet. In this case, each hibernaculum would have their own pathogen population and fungal individuals would rarely be exchanged between them. Thus, if the environment within the hibernacula is the main source of bat infection, we would expect to find differentiated populations (i.e., different pools of MLGs) of *P. destructans* across different hibernacula (expectation 1). To test this prediction, we collected the pathogen from nine closely situated hibernacula in a region of North-Eastern Germany. Second, we investigated the transmission pathway in great detail in a single intensively surveyed site (Eldena; one of nine sites used in the regional comparison above); that is, between hibernaculum walls (the suspected reservoir) and bats (the hosts). We tracked *P. destructans* MLGs from hibernaculum walls to bats through time and infer some important parameters relevant to this transmission (i.e., directionality, extent). If bats become infected from inside the hibernaculum, we expect the relative MLG abundances to remain mostly constant over consecutive winter seasons (expectation 2). That is because hibernacula would act as closed systems with little to no exchange of MLGs with other hibernacula and would support the long-term persistence of *P. destructans* spores in stable environmental conditions (as is the case in hibernacula). If bats indeed become infected from the walls of the hibernaculum, the MLG richness of *P. destructans* on the wall is expected to be greater than on the bats following the classical theory of transmission bottlenecks, whereby pathogen diversity is reduced on a host as not all MLGs successfully infect the host (expectation 3).The differences in MLG richness between walls and bats can also be used to quantify the strength of the bottleneck, and hence infer the quantity of spores transferred from walls to bats. Given the difficulty in obtaining large numbers of viable *P. destructans* spores from hibernacula walls (due to overall lower quantities of spores in the environment), we expect that the infection originates from a small number of spores transmitted from the walls to the bats (expectation 4).

**Figure 2.**
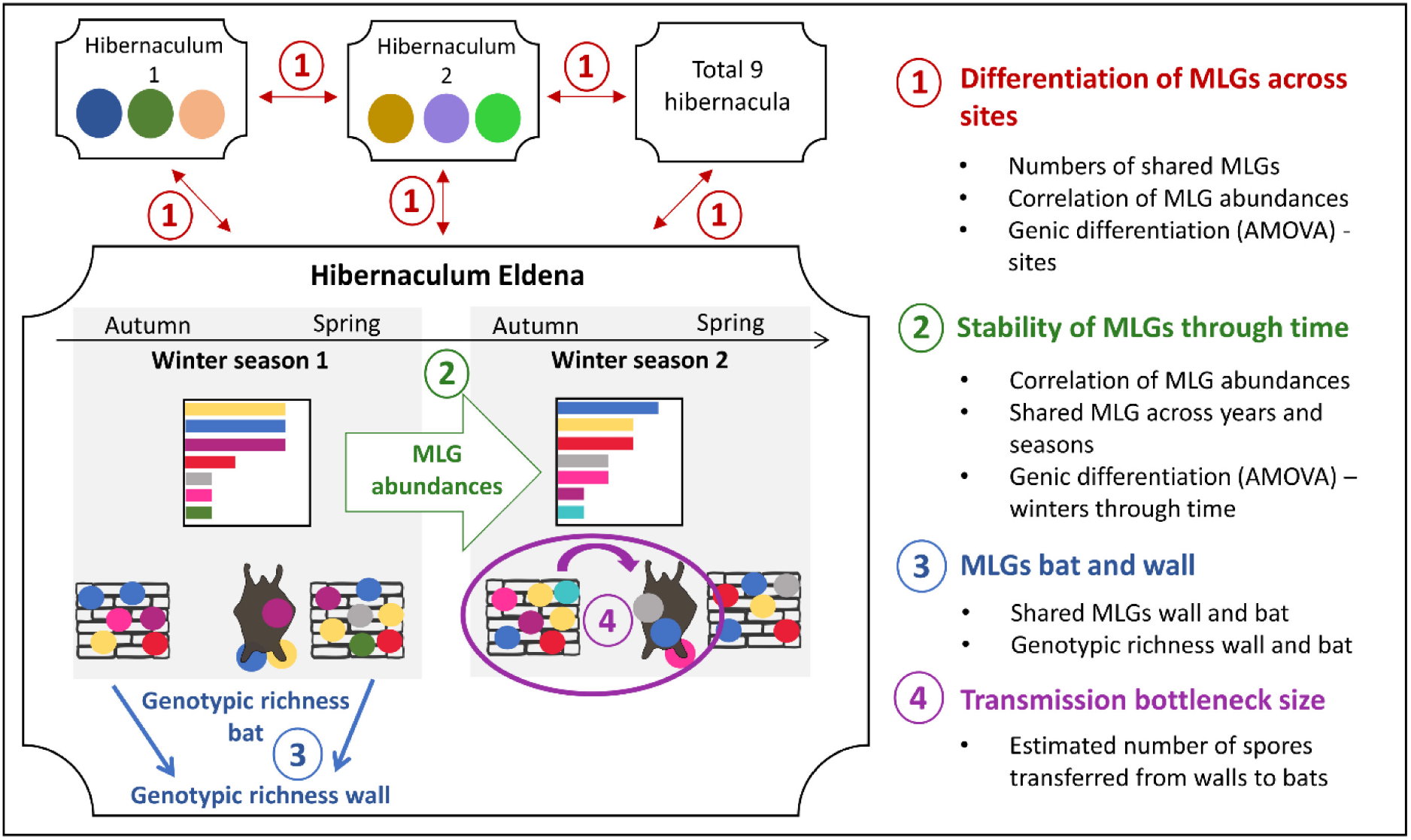
Summary of analyses used to address the four expectations concerning environmental infection of bats with *P. destructans*. Coloured dots and bars represent the identity of MLGs (quantity of MLGs is underrepresented for visual purposes). A reduced number of hibernacula and winter seasons are shown to illustrate the overall framework in a concise manner.

## Methods

### Sampling

Swab samples were collected from bats in nine bat hibernacula in the North-Eastern German federal state of Mecklenburg-Vorpommern, which is situated within the native range of *P. destructans*. These hibernacula are not used in summer by maternity colonies. Swab samples were collected from freely hanging and visibly infected bats (*Myotis myotis, M. nattereri, M. dasycneme*) between January and April. Bats were not handled for this sampling. Eight sites were surveyed only once or twice to obtain a snapshot of *P. destructans* populations to compare with the main study site, Eldena, for which the sampling was more extensive as explained below.

In the main study site, Eldena, both bats and walls were sampled across a period of four years (2015-2019). During bi-weekly visits to Eldena between 2015 and 2018, the number and species of bats present was documented (see supplementary material S1). The walls of the hibernaculum were also sampled by swabbing areas within 10 cm from where bats usually hung during hibernation (based on observations made during the bi-weekly bat counts). The surface of the wall swabbed was comparable to the area swabbed for the bats’ body parts. This wall swabbing was done over 4 consecutive years, twice a year; once in late April, to capture the time when most hibernating bats had left the site, and once in mid-October to sample just before the arrival of the majority of bats for the next swarming and hibernation season (see supplementary material S1).

### Culturing

Given that a swab sample, whether collected from a bat or a wall, often contains multiple MLGs (Dool et al., 2020), it is important to first isolate single spores before proceeding with culture and DNA extraction. To do so, each sample was cultured and upon germination of spores (visible under the microscope within 2-4 days after plating), individual spores were physically separated into fresh petri dishes to obtain single-spore isolates (SSI). All swab samples were cultured on dextrose-peptone-yeast agar (DPYA) following Vanderwolf et al. (2016). All SSI were sealed in petri dishes and stored upside down at 10°C for at least 6 weeks before they were harvested for DNA extraction. Depending on availability, in Eldena 1 to 3 (mean = 2.75, median = 3) and 1 to 5 (mean = 3.51, median = 5) SSI were isolated for bat and wall swabs respectively, while 1 to 4 (mean = 2.86, median = 3) SSI were isolated from the bat swabs obtained at other sites.

### Molecular analyses

In preparation for DNA extraction, fungal material was harvested from the SSI and dried (Vacuum centrifuge, V-aq, 30°C, 2.5 h) before homogenization in a tissue lyser (3x 15 sec. at 25 Hz) after 20 minutes at −80°C. The DNA was then extracted using a KingFisher Flex extraction robot (Thermo Scientific) and the MagMAX™ Plant DNA Isolation Kit (Thermo Scientific) utilizing magnetic-particle technology with the addition of 40 μM dithiothreitol (DDT) to lysis buffer A and 16.33 μM RNase A to lysis buffer B. To genotype isolates of *P. destructans*, 18 microsatellite markers (Drees et al., 2017b) were used in 4 PCR multiplexes as described in Dool et al. (2020). Genotyping was carried out using an ABI 3130 Genetic Analyser (Applied Biosystems). GeneMapper^®^ Software v.5 (Applied Biosystems) was used for fragment analysis.

### Data analysis

Most of our analyses use genotypic data based on the identification of MLGs which are defined by a distinct combination of alleles at the 18 microsatellite loci (Figure 2). To ensure the marker set was powerful enough to distinguish present MLGs, we calculated 1/probability of identity (P_ID_), which gives the theoretical number of different MLGs that can be distinguished (Waits et al., 2001). To determine the genotypic richness (diversity based on quantity of different MLGs), we used the measure of eMLGs which is the number of expected unique MLGs at the smallest shared sample size across several groups/populations based on rarefaction, as classically used to calculate allelic richness (Leberg, 2002). Its advantage is that it is independent of sample size and can therefore be used to compare the genotypic richness in groups of interest with differing sample sizes. The results for Eldena were obtained by using up to 3 isolates per swab for bats and 5 isolates per swab for walls. This was done to obtain large sample sizes to better estimate genotypic richness overall as well as at each sampling event. To ensure that patterns of genotypic richness were not influenced by differences in SSI per swab, we also analysed the data using exactly 3 SSI per swab independent of substrate, which showed the same patterns of genotypic richness (see supplementary material S4).

In addition to the analysis of genotypic patterns, we used AMOVA (analysis of molecular variance; using 1000 permutations for significance testing) based on allele frequencies. This method was used to test for genic (based on allele frequencies) differentiation between different groups of interest. All analyses were performed in R (version 4.0.2; R Team, 2019) using the packages poppr (version 2.8.6, Kamvar et al., 2015; Kamvar et al., 2014b) and vegan (version 2.5.6, Oksanen et al., 2019). The tidyverse collection of packages (Wickham et al., 2019) was used to improve ease and efficiency in analyses and the corrplot package (Wei & Simko, 2017) was used for visualization of correlations. The R-script used for analyses as well as the raw data are available as electronic supplementary material.

### Expectation 1: Geographic genotypic and genic differentiation – Are spores exchanged between bats during the summer?

If exchange of viable *P. destructans* was occurring bats in maternity colonies, i.e., during the summer and outside of the hibernacula, it would influence the patterns of genotypic and genic differentiation between hibernacula. We therefore described the patterns of genotypic and genic differentiation between hibernacula using two complementary methods. Firstly, we investigated genotypic differentiation across the nine sites by correlating MLG abundances between each pair of sites (Pearson product-moment correlation, p-values corrected using sequential Holm-Bonferroni method), with a strong positive correlation expected between sites with high rates of MLG exchange which would lead to reduced genotypic differentiation. Secondly, in addition to genotypic measures of richness and differentiation, we used AMOVA to determine if there was genic differentiation between the nine sites.

### Expectation 2: Stability of MLG through time and substrates (Eldena) – Do individual MLGs persist long-term?

If bats become infected from the walls of their hibernacula and little exchange of MLGs is happening across sites, hibernacula can be considered as closed systems for *P. destructans*. In this case we would expect a high stability of each MLGs’ abundance through time, with common MLGs in one year remaining common in the next. Therefore, the temporal stability of MLG composition in Eldena was studied across different times of the year (October & January-April) and different substrates (bat & wall) between 2015 and 2019. This was visualized by showing the presence of observed MLGs (excluding MLGs which only occurred at a single sampling event) at different sampling events as well as their respective overall abundance (based on an R script provided by Z.N. Kamvar, see Kamvar et al., 2014a). The stability of MLG abundances was evaluated by pairwise correlations between hibernation seasons (also referred to as “winter seasons” from now on; includes the wall swabs collected in October pooled with bat and wall swabs collected between January and April the next calendar year). For this we calculated a correlation matrix (pairwise correlations based on Pearson product-moment correlation, p-values corrected using sequential Holm-Bonferroni method) from the abundance of each MLG in Eldena (including unique MLGs) per winter season (2014/15 – 2018/19). This was done to determine whether common MLGs remained common or if there was substantial turn-over in the pathogen population across five winter seasons.

An AMOVA based on allele frequencies was performed to determine if genic differentiation existed across time (i.e., winter seasons).

### Expectation 3: Investigating genotypic and genic differentiation between bat and wall SSI in Eldena – Is there a transmission bottleneck?

We investigated the presence of environmental infection (i.e., infection from the walls of the hibernaculum) by studying the isolates/MLGs observed in samples obtained from bats (“bat isolates”/ “bat MLGs”) as well as from walls (“wall isolates”/ “wall MLGs”) in Eldena. If the walls of hibernacula are the source of bat infection, we expect to find genotypic signatures consistent with a transmission bottleneck, whereby the genotypic richness is reduced following infection from wall-to-bat as not all MLGs are transferred to the host population. To examine the existence of such a transmission bottleneck, we calculated eMLGs for bat and wall isolates to compare genotypic richness across substrates and years (subsampling and permutation of SSI were used to test for significance, see supplementary material S3.2). Furthermore, if frequent exchange of viable *P. destructans* spores was happening between walls and bats, we would expect to see a lack (or low amount) of both genic and genotypic differentiation between bats and walls SSIs. To test this, we correlated the observed MLG abundances found on bats and walls using Pearson product-moment correlation and used AMOVA (after clone-correction) based on allele frequencies to quantify genic differentiation between the two substrates.

### Expectation 4: Investigating the wall to bat transmission bottleneck in Eldena – How many spores are transferred per bat?

The difference in genotypic richness on the donor (walls – environmental reservoir) and receiver (bats – hosts) populations can be used to estimate the strength of the transmission bottleneck (i.e., number of spores passed) at the start of the hibernation season. To do so, we simulated bottlenecks of different strength (i.e., sample size; in steps of 1 from 2 to 3100) by subsampling the pool of SSI isolated from the walls (i.e., spores with known MLGs) in Eldena, mimicking the subset of spores passed from the walls to the bats (each winter season investigated separately). As previous studies have shown that only a fraction of spores is germinating, we subsequently randomly selected 17.5% of the previously subsampled SSI for further analyses (based on the mean germination rate for spores of *P. destructans* from the data in Fischer et al., 2020; mean = 17.5%). For each value of the bottleneck strength (i.e., sample size), we then calculated the genotypic richness (i.e., number of MLGs) observed in the subsampled dataset. We then compared the results from all simulations to infer the bottleneck size with the average (over 1,000 runs) that best matched the observed genotypic richness in bats in Eldena (supplementary material S3.1). Although we isolated and genotyped several SSI per sample (see above), it is likely that we did not capture all the MLGs that were present on the samples we analysed. Hence, to get more precise results, we also calculated the number of transferred spores obtained when using a predicted number of MLGs for bat samples we analysed in addition to the observed number. This predicted number of unique MLGs for bat samples was obtained via a Bayesian estimator classically used to estimate population sizes based on a single sampling session (Petit & Valiere, 2006; Puechmaille & Petit, 2007; supplementary material S3.1).

To obtain an average number of spores passed from the wall reservoir to each single bat, we simply divided the determined matching bottleneck size (i.e., number of spores transferred to the entire sampled population of bats) by the number of body parts that were sampled (i.e., the number of samples) and multiplied by six, the number of body parts that are commonly infected by *P. destructans* (i.e., left/right ear, left/right wing, nose and uropatagium).

## Results

### Geographic genotypic and genic differentiation (expectation 1)

For Eldena, (the focal site), 287 bat swabs and 78 wall swabs were processed over the course of 5 winters (2014/15 until 2018/19), resulting in a total of 1062 *P. destructans* single spore isolates (SSI). The number of swabs collected from the 8 other sites was at least 10 and resulted in 27 to 85 SSI per site, for a total of 435 SSI (Table 1). All presented SSI were genotyped at 18 microsatellite loci with an overall level of missing data < 1% (across all sites and loci). The P_ID_ across sites was 6.63 *10^−7,^ indicating that more than 1.5 million different MLGs could theoretically be distinguished by our marker set, clearly exceeding the obtained number of MLGs (see supplementary material S2.1 for P_ID_ at each site).

**Table 1.** Information on sample sizes, genotypic richness, and abundance of shared MLGs across sites. Distance in km refers to the distance from each hibernaculum to Eldena. N_Swabs_ refers to the number of swab samples collected and cultured with N_SSI_ giving the number of obtained single spore isolates and N_MLG_ giving the total number of unique multi-locus genotypes per site. Shared MLGs are given as all MLGs shared with any other site; the number of MLGs shared with Eldena are given in brackets.

| Hibernaculum | Distance [km] | $N_{Swabs}$ | $N_{SSI}$ | $N_{MLG}$ | Shared MLG |
| --- | --- | --- | --- | --- | --- |
| Eldena | - | 365 | 1062 | 149 | 13 |
| Wolgast | 21 | 22 | 63 | 18 | 3 (2) |
| Anklam | 30 | 12 | 33 | 16 | 3 (0) |
| Richtenberg | 39 | 12 | 35 | 7 | 5 (4) |
| Altentreptow | 46 | 27 | 76 | 31 | 8 (5) |
| Bad Sülze | 52 | 10 | 31 | 11 | 2 (1) |
| Strasburg | 67 | 34 | 85 | 28 | 8 (6) |
| Burg Stargard | 67 | 10 | 27 | 18 | 3 (1) |
| Neustrelitz | 83 | 32 | 85 | 32 | 9 (1) |

Each site had a nearly unique collection of MLGs with 92 % of MLGs not shared between sites (278 MLGs in total, of which only 22 were shared between two or more sites; see supplementary material S2.2). Of the 149 MLGs observed in Eldena between 2015 and 2019, only 8.7% were shared with other sites (13 MLGs). The abundance of each MLG showed low and non-significant correlations between sites, ranging from −0.09 to +0.12 (all p > 0.05; see supplementary material S2.3). The correlation of MLG abundance observed in Eldena with Strasburg and Neustrelitz (which had the greatest numbers of SSI apart from Eldena; see Table 1) was −0.04 and −0.09, respectively. Even the MLG abundances pooled for the eight sites excluding Eldena (N_SSI_ = 435) were not-significantly correlated with the MLG abundance in Eldena (all years pooled, N_SSI_ = 1062; *df* = 356, r= −0.05, *p* = 0.80).

Furthermore, AMOVA based on allele frequencies at the 18 microsatellites detected significant genic differentiation between sites (*df* = 309, 4.1 % of variance explained by sites, *p* = 0.001).

### Stability of MLG through time and substrates (Eldena; expectation 2)

We found a great number of MLGs in Eldena shared across different sampling events and years (Figure 4). To investigate the turnover of MLGs and their abundance over time in Eldena in more detail, we calculated a correlation matrix based on the abundance of each MLG (including unique MLGs) for pairwise winter seasons (5 winters: 10 pairwise comparisons). We found high and significant correlations between all pairs, ranging from 0.64 to 0.76 (Pearson product-moment correlation, p < 0.001 for all pairs; Figure 3). The correlation between the first and last sampled winters (2014/15 & 2018/19) was 0.67. In agreement with the high stability of MLG abundances in Eldena, the genic differentiation (based on allele frequencies) was not significant between winters (AMOVA, *df* = 285, 0 % variance explained by winter seasons, *p* = 0.987).

**Figure 3.**
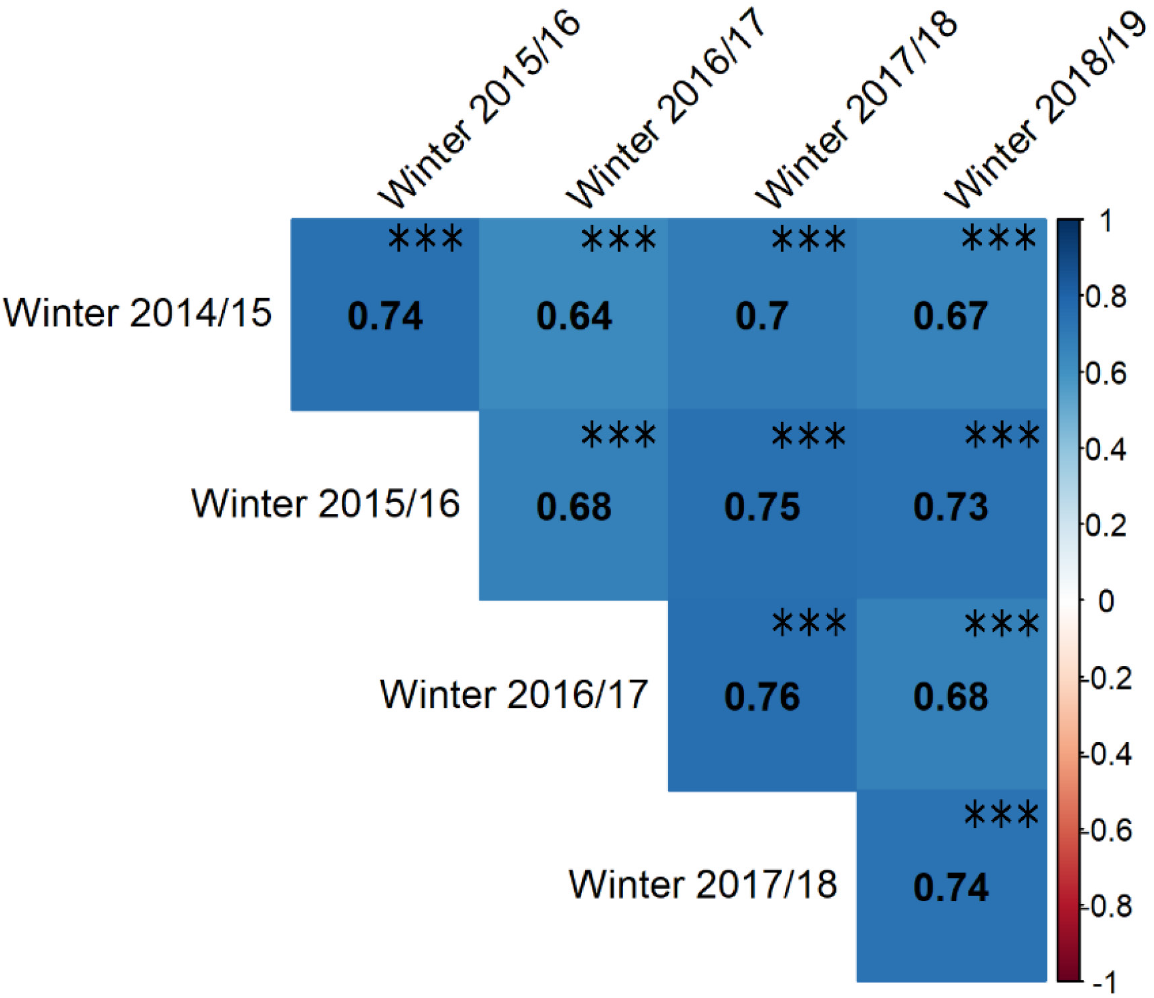
Correlation of MLG abundances (from bats and walls) across five different winter seasons in Eldena (winter 2014/15 until 2018/19). For both, values and colour represent Pearson product-moments correlation indices while stars indicate level of significance after sequential Bonferroni-Holt correction (***: p < 0.001).

**Figure 4.**
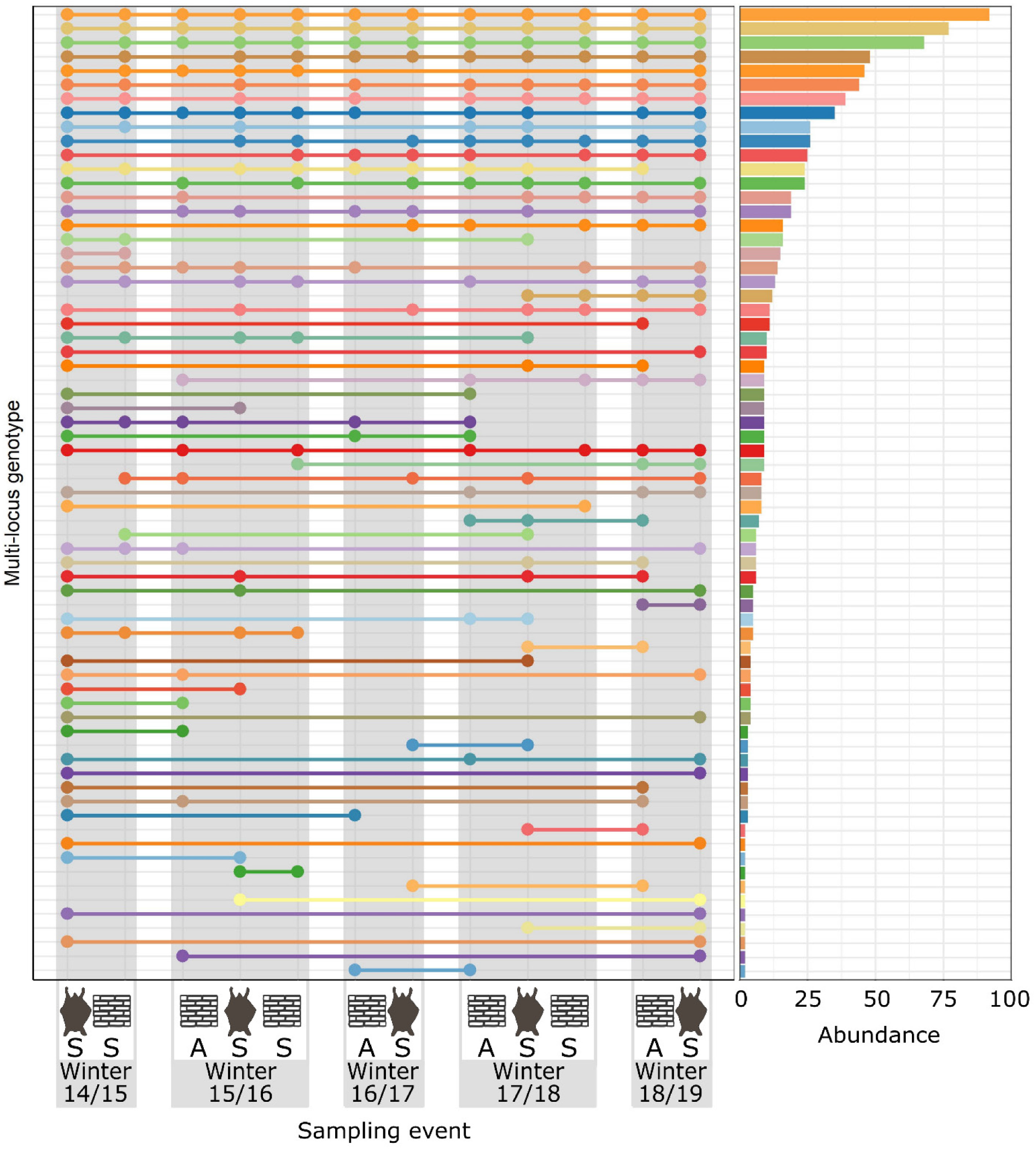
On the left, the observation of MLGs (y axis) across different sampling events (y-axis) is shown, whereby substrates are indicated by bat and wall symbols, sampling times by letters (A = autumn, S = spring) and winter seasons by grouping of sampling events in grey bars. A dot at a sampling event shows that an MLG was observed (absence of a dot indicates that the MLG was not observed at this event). On the right side, the overall abundance of each MLG is shown. Only MLGs observed at more than one sampling event are shown.

### Investigating genotypic and genic differentiation between bat and wall SSI in Eldena (expectation 3)

The sampling of Eldena spanned a period of 4 years with sampling in regular intervals (walls in October and April & bats between January to April) between 2015 and 2019 which yielded 788 bat SSIs and 274 wall SSIs. Considering an equal sample size, the smallest shared sample size of 274 SSI, genotypic richness of *P. destructans* was higher on samples collected from walls than from bats (eMLG_WALLS_ = 85, eMLG_BATS_ = 76.5; permutation test, p < 0.001, see supplementary material S3.2). Per winter season, eMLG values of wall samples (pooled per winter, i.e., October and April) were higher than bat samples (with the exception of winter 2014/15; Table 2).

**Table 2.** Information on sampling success (*N_Swabs_* = Number of cultured swab samples, *N_SSI_* = Number of single spore isolates) and genotypic richness (*N_MLG_* = Number of unique MLGs, *N_eMLG_* = Number of expected MLGs at equal sample size) in winters in 5 years and from different substrates (bat/wall) for Eldena.

| Winter & substrate | $N_{Swabs}$ | $N_{SSI}$ | $N_{MLG}$ | $eMLG$ |
| --- | --- | --- | --- | --- |
| 2014/15 bats | 140 | 391 | 74 | 16.9 |
| 2014/15 wall<sup>a</sup> | 17 | 72 | 33 | 16.5 |
| 2015/16 bats | 31 | 80 | 34 | 15.3 |
| 2015/16 wall | 16 | 39 | 29 | 18.2 |
| 2016/17 bats | 10 | 26 | 16 | 14.4 |
| 2016/17 wall<sup>b</sup> | 6 | 22 | 17 | 17.0 |
| 2017/18 bats | 53 | 149 | 37 | 14.5 |
| 2017/18 wall | 24 | 89 | 37 | 15.7 |
| 2018/19 bats | 53 | 142 | 47 | 15.7 |
| 2018/19 wall<sup>b</sup> | 15 | 52 | 32 | 16.9 |

<sup>a</sup> contains isolates from wall sampling in April only.
<sup>b</sup> contains isolates from wall sampling in October only.

We detected considerable numbers of *P. destructans* MLGs shared between bats and walls in Eldena and a high quantity of shared MLGs across the five different winter seasons (Figure 4). When data from the five winter seasons were pooled together, the Pearson correlation of MLG abundances sampled from bats and those sampled from walls was high (r = 0.88, *p* < 0.001). There was no significant genic differentiation between the sampled substrates (walls and bats, AMOVA: *df* = 311, 0% variance explained by substrate, *p* = 0.974)

### Investigating the wall to bat transmission bottleneck in Eldena (expectation 4)

Based on the subsampling of SSI (i.e., genotyped spores) isolated from the walls (which mimics the bottleneck occurring when bats get infected from the walls), the germination rate and the sampling scheme/intensity, we estimated from the observed number of MLGs that between 49 and 104 spores were transferred from walls to each bat at the start of hibernation. When considering the predicted number of MLGs, the estimates were in the range of 48 – 530 spores/bat (supplementary material S3.1). The estimate obtained for the winter season with the most intense sampling (i.e., the winter 2014/15) is between 100 and 129 spores transferred from the walls to each individual bat.

## Discussion

### Geographic genotypic and genic differentiation (expectation 1)

While the majority of observed *P. destructans* multi-locus genotypes (MLGs) were restricted to single hibernacula, we did observe a small fraction of MLGs present at more than one site. While *P. destructans* cannot grow on active bats (Verant et al., 2012), Campbell et al. (2020) found that spores could survive at elevated temperatures for several weeks, suggesting that *P. destructans* can remain viable on active bats for periods that are sufficient for bats to move it from one site to another. The successful movement of viable spores should primarily depend on the number of inter-site bat movements and the number of viable spores on the bat when such movements occur. Hence the late hibernation/spring transition period is the most favourable time for successful transfer of spores between hibernacula. Indeed, the fungal load of *P. destructans* on bats is at its highest in spring when the bats are in their final stage of hibernation (Langwig et al., 2015a; Puechmaille et al., 2011). At this time, bats typically start moving between sites – from hibernacula to summer roosts – possibly transiting via other hibernacula in between. Although bats groom off most of the visible fungal material upon waking from hibernation, some spores and fungal hyphae are still present on their skin (Puechmaille et al., 2011). It remains unclear whether only few spores manage to be transferred or survive between sites or, if more than the 8% of shared MLGs between sites observed in this study are transferred but are then unable to survive and/or successfully infect bats. Although beyond the scope of the current study, factors like local adaptation to environmental conditions as well as intra- and interspecific competition might play a role that remains to be investigated (Lilley et al., 2018).

Our population genetic data, suggesting low exchange rates of MLGs between hibernacula, might at first seem contradictory with the rapid colonisation of the fungus in North America during the last decade (www.whitenosesyndrome.org). However, the geneflow between sites already occupied by the fungus and the colonisation of novel sites are different processes. The colonisation of a site previously not harbouring *P. destructans* requires that the pathogen has the ability to cope with local abiotic environmental conditions and local biota (possible inter-specific competition). In addition to these factors, MLGs dispersing between sites with already existing *P. destructans* populations also need to cope with possible competition between MLGs (intra-specific competition). In addition, colonisation is likely aided by more frequent arousals of heavily infected bats experiencing the novel pathogen in North-America (Reeder et al., 2012) which might further facilitate the spread of the pathogen between sites. Further work is needed to differentiate the role of preceding (i.e., historic) colonisation of MLGs across sites in Eurasia from contemporary geneflow.

Our data suggest that if successful geneflow between hibernation sites with established *P. destructans* populations is happening, it is a relatively rare event (compared to in-situ recruitment), which does not significantly modify the patterns of genotypic richness (at least in the species native range at the time scale investigated herein).

### Plausibility of *P. destructans* persistence on bats over summer

*P. destructans* is highly temperature sensitive with optimal growth between 12 and 15 °C and complete cessation of growth above 20°C (Verant et al., 2012). Hence growth is not possible on active bats given that their body temperature is around 37°C when active and the ambient temperature is known to reach temperatures of 40°C during the day in maternity colonies, where females and juveniles spend several months in summer (e.g. Zahn, 1999 for *M. myotis*). Fungal spores are known to be more durable than fungal hyphae, but even *P. destructans* spore viability decreases to a few percent after only 15–20 days at 37°C (Campbell et al., 2020; Veselská et al., 2020), suggesting that the fungus is unlikely to remain viable on bats through the active summer period. Screening results for *P. destructans* on bats during the summer is consistent with this prediction with only a small fraction of individuals tested carrying DNA of the fungus (e.g. Ballmann et al., 2017; Carpenter et al., 2016; Huebschman et al., 2019; Langwig et al., 2015a). Given that the large majority of studies on summer prevalence are based on qPCR data, it remains unclear what fraction of these infrequent qPCR positives relate to viable spores. Results from two studies attempting to cultivate the fungus from the wings of active bats sampled during the summer (June–August), provided evidence that viable *P. destructans* was rarely encountered (2 positives out of 117 samples cultured; Ballmann et al., 2017; Dobony et al., 2011). One positive was a juvenile *Myotis lucifugus* (sampled on 19/08) while the second one was an adult female *Myotis lucifugus* (*s*ampled on 15/08; Ballmann et al., 2017; Dobony et al., 2011). For the juvenile, carryover of the infection from the previous winter can be excluded. For the adult, a new infection is also most likely given that the individual was captured during the early swarming season (Thomas et al., 1979) in a hibernaculum where *P. destructans* is known to occur, including on wall surfaces (Ballmann et al., 2017). Although there are technical challenges to isolating viable *P. destructans* from individuals that carry a small number of spores, all studies conducted so far suggest that *P. destructans* (hyphae as well as spores) infecting bats during the hibernation season is not able to remain viable on bats through the active summer period, and unable to cause infection in autumn. In other words, a yearly contamination is needed for the pathogen to infect its host and complete its life cycle.

The yearly re-infection of bats is completely concordant with a limited pathogen exchange between different hibernacula. It has been shown that susceptible, healthy bats can become infected by others carrying the fungus during hibernation (Lorch et al., 2011). Considering that temperate bats also hang close to each other and often form dense clusters in maternity roosts, frequent exchange of pathogens like *P. destructans* spores on fur/skin would be very likely (e.g. Webber & Willis, 2016). Bats from the hibernacula sampled for this study use the same maternity roost in summer (Steffens et al., 2007; bat ringing centre Dresden pers. comm., data collected between 1981 – 2019). We would therefore expect a mostly homogenous pathogen population with weak spatial structure of *P. destructans* across these sites if the fungus remained viable and infectious throughout the summer on bats. On the contrary, we observed striking differences in genotypic distribution between sites with 92% of MLGs found at only one site, thereby confirming expectation 1 for wall-to-bat infection within the hibernacula. We conclude that the large majority of bats returning to hibernacula have cleared the infection over summer and that re-infection stems from within the hibernacula. The differentiation between sites might be further exacerbated by the widespread faithfulness of bats to their hibernacula within a winter season (Reusch et al., 2019; Steffens et al., 2007). The timing of infection is most likely during the autumn swarming season as previously hypothesised (Puechmaille et al., 2011), when contact between walls and bats is exacerbated. This period only briefly precedes the onset of hibernation, allowing spores to remain viable (up to 15 days at 37°C, Campbell et al., 2020) and cause an active infection during hibernation (when bat body temperature is reduced) shortly after.

### Investigating genotypic and genic differentiation between bat and wall SSI in Eldena (expectation 3)

Temperate zone bat hibernacula are characterized by cool, constant temperatures (< 20°C all year) with little fluctuation between day and night as well as high humidity (e.g. Perry, 2013). These conditions have been shown to allow the persistence of viable *P. destructans* spores on abiotic substrate for at least 2 years (Fischer et al., 2020). In addition to walls being likely abiotic reservoirs, sediments as well as bat guano have been suggested to be possible sources of *P. destructans* environmental infection (Lorch et al., 2013; Urbina et al., 2020). However, biotic interactions between fungi and other microorganisms are most likely stronger in sediments/guano compared to hibernacula walls, making sediments/guano a challenging environment for *P. destructans* to survive in and grow (Zhang et al., 2014, Wilson et al., 2017, Urbina et al., 2021). Furthermore, physical contact between bats and guano/sediments is a rare event in most bat species, suggesting these potential reservoirs are unlikely to play an important role in *Pd* infection. In contrast, contact of bats with the walls of hibernacula occurs often as bats land and crawl on walls during swarming and hibernation and contact is also often observed during grooming in arousal bouts from hibernation. From this, we conclude, that sediments and guano are unlikely to play a significant role as pathogen reservoirs from which bats could become infected.

Following the classical theory of transmission bottlenecks, not all multi-locus genotypes (MLGs) successfully transfer from a source population to a host. This results in the pathogen population on the host including only a subset of the population available on the source with reduced richness. Here, we confirmed expectation 3 of wall-to-bat infection by showing that bats harbour lower genotypic richness for *P. destructans* than walls, with the number of eMLG being 10% lower on bats than on walls (MLGs pooled across all sampling events). From this we can conclude that walls are the source population from which bats become infected every year. This conclusion is consistent with recent studies finding similar fungal and bacterial assemblages on bats wings and their hibernacula walls, emphasising the strong effect of site on the bat skin microbiome, largely consisting of site-specific microbiota (Ange-Stark et al., 2019; Avena et al., 2016). These findings suggest frequent exchange of microbial material with a potential source-sink relationship of microorganisms between bats (the potential sink) and their environment (the potential source; directionality of source-sink likely depending on the microbial species, Grisnik et al., 2020). In the case of *P. destructans*, the situation is more complex as the environmental reservoir is replenished from the bats at the end of the hibernation season.

### Investigating the wall to bat transmission bottleneck in Eldena (size of bottleneck; expectation 4)

Understanding the quantity of spores with which bats are infected is an important aspect in the disease dynamics. For example, considering that disease severity and response might be dependent on inoculum load, using appropriate fungal loads would be essential to obtain accurate results in studies based on artificial infection of hosts. We estimated the spore load (the number of spores passed from the walls to one bat at the start of the season), to be between roughly 50 and 500 spores. This finding is consistent with the expectation that only a limited amount of spores is transferred from walls to bats during infection in autumn based on the difficulty in obtaining viable fungal material from the walls of hibernacula (unpublished data); it appears unlikely that bats come in contact with large quantities of spores. While this represents a rough approximation of transferred spores that could be estimated more accurately in subsequent studies, we believe that the order of magnitude of our findings (i.e., approximately several hundred spores per bat) is likely correct. Hence, initial pathogen load is remarkably lower than the inoculum sizes used in most studies artificially infecting bats with *P. destructans*, which was typically around 500.000 spores per wing (e.g. Lorch et al., 2011, Warnecke et al., 2012). A study by Johnson et al. (2014) found that the inoculation of *Myotis lucifugus* bats with 500 spores of *P. destructans* reduced the bats survival odds significantly which was not observed for higher inoculum sizes (5,000 – 50,000 and 500,000 spores tested; potential explanations for this non-intuitive result are discussed by the authors). Our results suggest that future studies aiming at elucidating physiological or behavioural aspects of infection in hosts in an experimental setup use an adjusted (i.e. lower) inoculum size (or ideally several inoculum sizes) in order to more closely match the apparent situation for wild bats naturally infected by *P. destructans*.

### Stability of MLG through time and substrates (Eldena) – Constancy of MLGs over time (expectation 2)

We found a high number of MLGs in Eldena to be shared across winters and substrates, with the most common MLGs being shared (in time and space) most often. This was expected, given that hibernacula can be seen as predominantly closed systems (i.e., no significant emigration, immigration or extinction) with a very stable environment. In simplified terms, we expect bats to become infected with the fungus from the walls of hibernacula in autumn. Over the winter, the fungus then reproduces clonally on bats by producing large amounts of spores. These spores will then be shed into the environment in spring when the pathogen load is highest. Some spores will survive on the walls of hibernacula through the summer and will be able to infect bats again when they return in autumn. In such a system we expect the relative MLG abundances to remain fairly constant: all other things being equal, the chances for an MLG to infect a bat in year n+1 will be proportional to its MLG abundance on the walls in year n. To test this, we examined the correlation of MLG abundances across years in Eldena which revealed a strong (and highly significant) correlation between each winter, even across multiple years.

With MLGs mainly specific to individual hibernacula and a yearly occurring transmission bottleneck (facilitating genetic drift) one question which remains unanswered is how the genotypic richness is maintained in the long-term within this mostly closed system. One possible factor is the occurrence of gene flow; although rare as seen above, it nevertheless possibly contributes to bringing in new MLGs and alleles. A second factor likely buffering the impact of differential success of MLGs per year (due to the transmission bottleneck) is the long-term survival of spores in the environmental reservoir demonstrated by Fischer et al. (2020). Indeed, MLGs that are not ‘picked up’ from the wall by bats in one year can remain on the wall and have further opportunities to infect bats in the subsequent years. This could stabilize MLG abundances as well as genotypic richness through time. A third and non -mutually exclusive possibility for genotypic richness to be maintained in the long-term could be the presence of sexual reproduction which would introduce novel MLGs. While sexual reproduction has never been observed in *P. destructans*, the presence of both mating types in hibernacula in the native range (Palmer et al., 2014) as well as the relatively even distribution of both mating types (frequency of 0.69:1 for MAT1_1/ and MAT1_2 across 1099 SSI from Eldena, unpublished data) suggest it might occur there, though likely less frequently than asexual reproduction (Dool et al., 2020). Additionally, spores resulting from sexual reproduction are often more durable than clonal spores (see e.g., Gersuk et al., 2006; O’Gorman et al., 2009 for *Aspergillus fumigatus)* which could further drive the long-term persistence of *P. destructans* in hibernacula environments. It would be very informative for future studies to investigate these aspects in greater detail; i.e., the role (and interplay) of gene flow, long-term survival and sexual reproduction in the maintenance of genotypic richness in *P. destructans*.

### Population genetics as a tool for understanding infectious disease dynamics

With an increasing human population as well as travel and the trade of goods across the world, the occurrence of novel pathogens and introductions of pathogens beyond their natural range are expected to increase. Therefore, understanding the processes involved in pathogen emergence and infection of hosts is critical to reduce the threat and manage outbreaks. Population genetics are an important part of the toolbox needed to reach this goal.

So far, population genetics have already been used to answer a broad range of questions related to pathogen biology (including parasitology). Examples of this include the elucidation of clonality in pathogens (Hill et al., 1995; Morehouse et al., 2003) and understanding the spatial structure and diversity of a pathogen in a geographical range (e.g. Anderson et al., 2000; Mekonnen et al., 2020) which can further be used to identify the source population of a pathogen’s introduction (e.g. Jarman et al., 2019). These types of studies investigate patterns of large (evolutionary) time frames, which are a valid and important part of the framework to understanding wildlife diseases (e.g. Vander Wal et al., 2014). Population genetics may also be used to understand disease in the context of short time frames such as the description of pathogen life cycles (including the study of alternative hosts or reservoirs). While such studies on pathogen reservoirs have been frequently conducted with regard to human or livestock pathogens (e.g., Dubey et al., 2020; Venkatesan & Rasgon, 2010) they have scarcely been conducted for wildlife pathogens. However, the knowledge on pathogen lifecycles and reservoirs is indispensable if we are to understand wildlife diseases, develop possible management strategies or indeed avoid the further spread of pathogens (e.g., from caving equipment in the case of *P. destructans;* Zhelyazkova et al., 2020; Zhelyazkova et al., 2019). Our study provides evidence that population genetic approaches provide an invaluable and resource-efficient tool to elucidate cryptic infection pathways, including in the field of emerging infectious diseases of wildlife where resources are often limited.

### Implications for conservation strategies

Traditionally, most management strategies to deal with emerging infectious diseases have focussed on targeting the host (see e.g. Langwig et al., 2015b). Among these are the controversial culling of infected hosts (see e.g. the culling of badgers in England to curb bovine Tuberculosis; Ham et al., 2019) as well as inhibition of the pathogen growth on hosts (by applying natural or chemical antifungals to hosts, e.g. Boire et al., 2016). Our result, providing clear evidence that the walls act as an environmental reservoir from which bats become infected with *P. destructans*, suggest that it is probably more feasible and effective to target the pathogen (instead of the host) to reduce fungal load in the environment and hence prevent yearly re-infection of the bat hosts.

If we consider White-Nose Disease dynamics, reducing the number of spores on the walls of hibernacula could lead to fewer bats becoming infected in the following hibernation seasons. With fewer infected bats in winter, the fungus would reproduce less on bats, and less spores would be shed on to the walls in spring. This could then lead to fewer infections of bats in the following year. Therefore, significantly reducing the amount of spores in the environment would be a straightforward way to break the cycle of re-infections, which can bring with it a long-term effect which would need to be repeated fewer times compared to methods targeting the host. The concept of reducing *P. destructans* load from hibernacula environments is supported by the work from Hoyt et al. (2020) who predicted that a reduction of fungal load to 20% of its current load over the summer could result in the stabilization of bat populations in North America.

Taken together, our study highlights that population genetic approaches can provide critical knowledge on pathogen transmission dynamics; knowledge that is critically needed to adjust management strategies, and that more studies are needed to focus on developing management strategies targeting the environmental reservoir of pathogens.

## Acknowledgments

We kindly thank Jens Berg, Anne Petzold, Holger Schütt, Dirk Karoske, Thorsten Blohm and Axel Griesau of NABU Mecklenburg Vorpommern bat conservation group for facilitating access to the sampling sites and Dagmar Brockmann of the bat ringing centre Dresden for sharing information on bat movements between the studied hibernacula. Furthermore, we thank Ina Römer, Silke Fregin, Ruth-Marie Stecker and Marcus Fritze for their help relating to labwork and sample collection.

## Data accessibility

The dataset supporting this article will be uploaded to Dryad upon acceptance of the article.

## Authors’ contributions

S.J.P conceived the study. N.M.F. and S.J.P. designed the study; S.J.P. acquired funding and supervised the project; S.J.P and G.K. administered the project. N.M.F. (40%), A.A. (25%), S.R. (25%), S.E.D (5%) and S.J.P. (5%) carried out the laboratory analyses; N.M.F. performed data analyses. N.M.F. and S.J.P. interpreted the results and wrote the original manuscript. All authors critically discussed the results and edited the manuscript, approved its final version, and agree to be held accountable for the content therein.

## Supplemental Information

### 1. Counts of bats in Eldena

Eldena was visited biweekly from July 2015 to July 2018 to establish the typical seasonality of bat presence and absence (Figure S1). Previous visits (less regular, hence not reported) were used to elucidate the best timing of sampling from bats and walls in Eldena. Most bats were sampled during March, when visible growth of *P. destructans* was greatest and most bats were present in the site. Samples from the walls were collected in April and October just before and after hibernation when most bats were absent.

**Figure S1.**
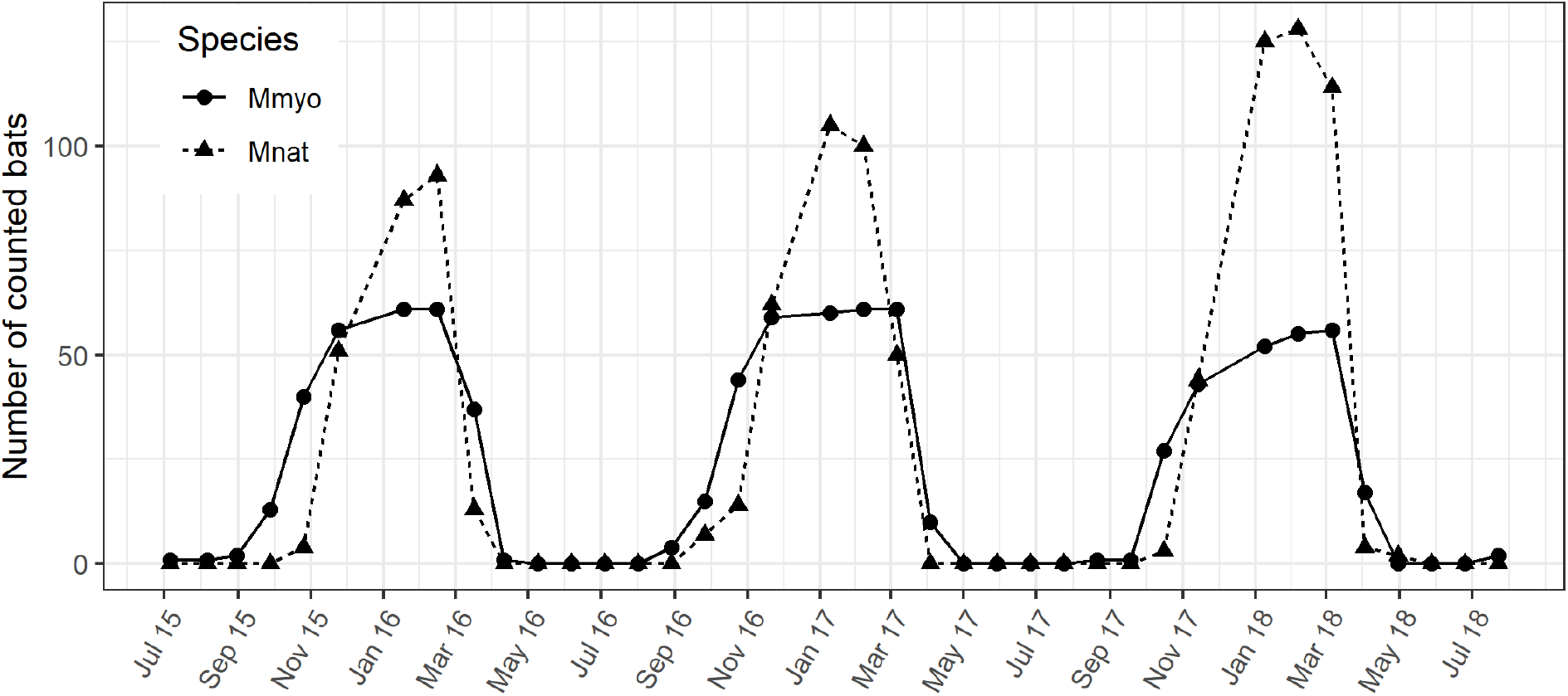
Number of *Myotis myotis* (dots) and *Myotis nattereri* (triangles) bats counted in Eldena every 14 days for 3 years. Counts data are shown for both species with symbols; the lines are only used to help visualise the patterns of changes.

### 2. Additional analyses on differentiation across hibernacula

If most bats lose viable fungal material (i.e., spores) over the summer, exchange of viable *P. destructans* MLGs between bats meeting in maternity colonies is expected to be rare. To investigate if this is the case, we compared the MLG populations from closely situated hibernacula. Bats from the sampled hibernacula meet in the same maternity colonies in summer (Steffens et al., 2007). Therefore, if bats indeed purge infection over summer, we expect to find differentiated MLG populations of *P. destructans* across the studied hibernacula.

All the studied hibernacula are underground human-made structures (i.e. cellars and bunkers). As it is often quite difficult to determine if close-by sites (e.g., part of the same bunker system) are connected we considered all hibernacula in the same town to be part of the same hibernacula cluster (simply referred to as hibernacula or sites).

#### 2.1 Probability if Identity (P_ID_) across sites

The P_ID_ gives the number of MLGs which can theoretically be distinguished using a specific marker set. This information is helpful in determining the reliability of MLG-identification across and between the studied hibernacula. The overall P_ID_ across sites was 6.63 *10^−7^ (1/PID > 1.5 million). Considering, that allelic diversity per locus was different across sites it was also important to estimate the P_ID_ for each of the sites to ensure that it was possible to determine reliable MLG identities for each one (used to find matches across the dataset to determine, e.g., the number of shared MLGs across sites). The number of 1/P_ID_ giving an estimate of distinguishable MLGs by far extends the number of observed MLGs (Table S1).

**Table S1.**
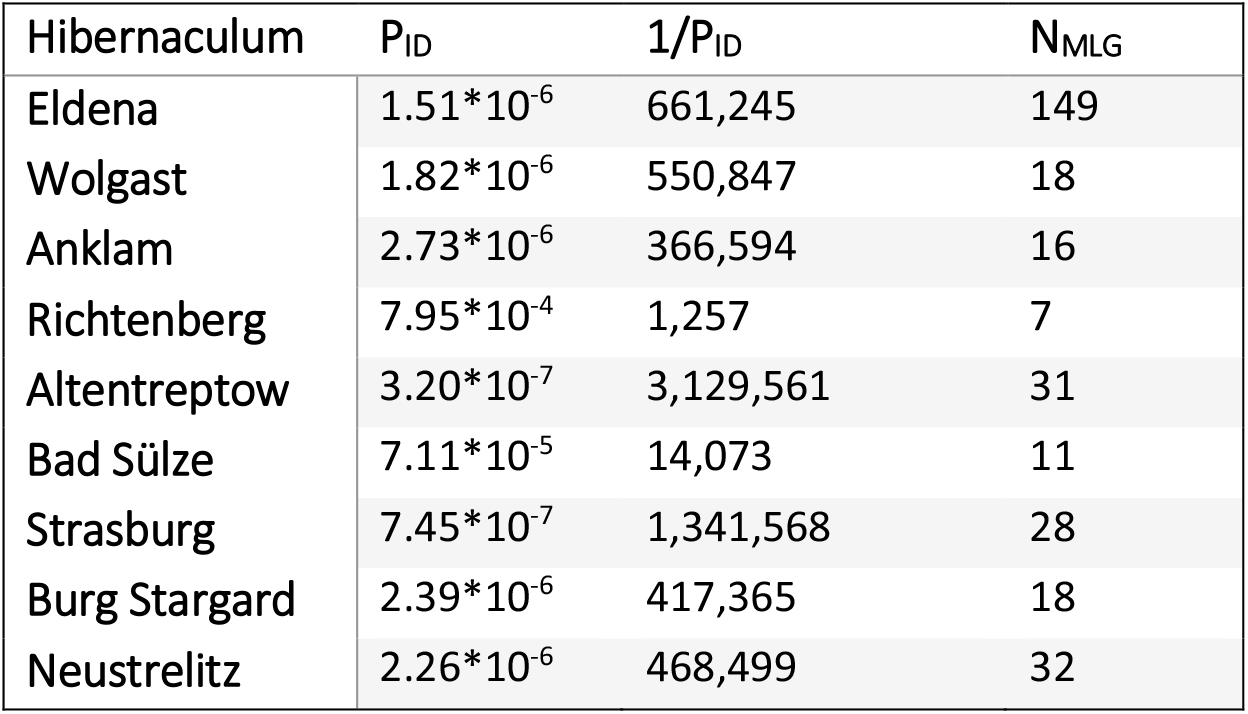
Probability of identity (P_ID_) and 1/ P_ID_ as well as the observed number of unique MLGs of *P. destructans* for the studied hibernacula

#### 2.2 MLGs shared across sites

Of all observed *P. destructans* MLGs, 92% were unique to sites with only a total of 22 MLGs shared between at least two sites (Figure S2). The most complete sampling was obtained for the hibernaculum Eldena, where 149 different MLGs were observed.

**Figure S2.**
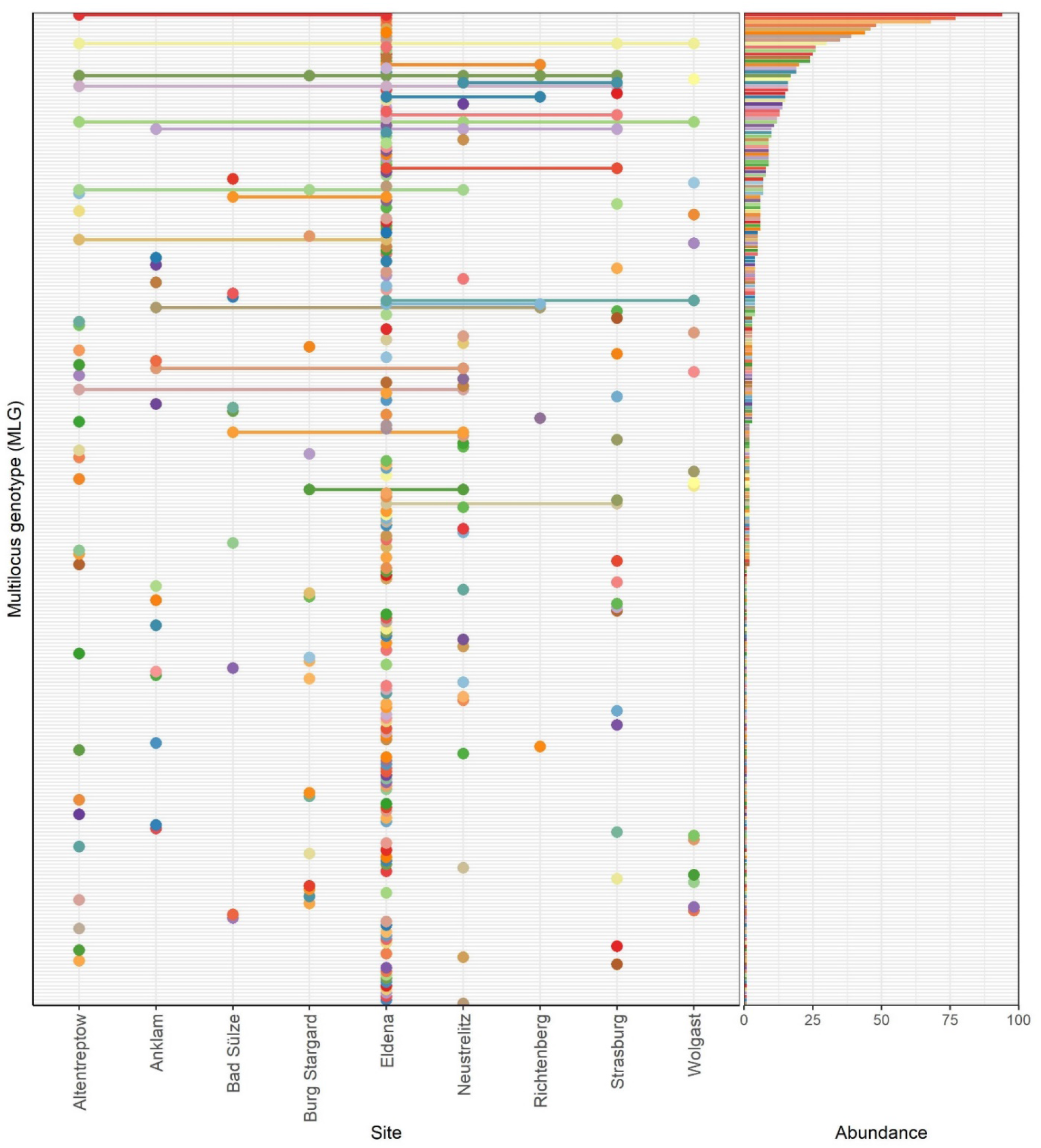
Observation of MLGs (y axis) across different hibernacula (x axis on left side). The left side of the graph shows the hibernaculum an MLG was observed at indicated by a dot (absence of dot shows that the MLG was not observed at this hibernaculum). On the right side, the overall abundance of each MLG across all sites is shown.

#### 2.3 Correlation of MLG abundances across sites

Based on MLG distributions in Eldena (see main text, Figure. 4) we know that some MLGs are more common than others, which is consistent across substrates (bat/wall) and time. MLGs which are common in one year are likely to be common in the following years. Therefore, if there was much exchange of MLGs across sites, likewise, we would expect that common MLGs would be more likely to be exchanged than rare ones leading to a high correlation of MLG abundances across sites (possibly slightly lower but comparable with the high correlation of MLG abundances between winters in Eldena which ranged from 0.64 to 0.76).

This was not the case with pairwise correlations (Pearson product-moment correlation) never exceeding 0.12 (Figure S3, *p* > 0.05 for all 72 pairs following sequential Bonferroni-Holm correction for multiple testing).

**Figure S3.**
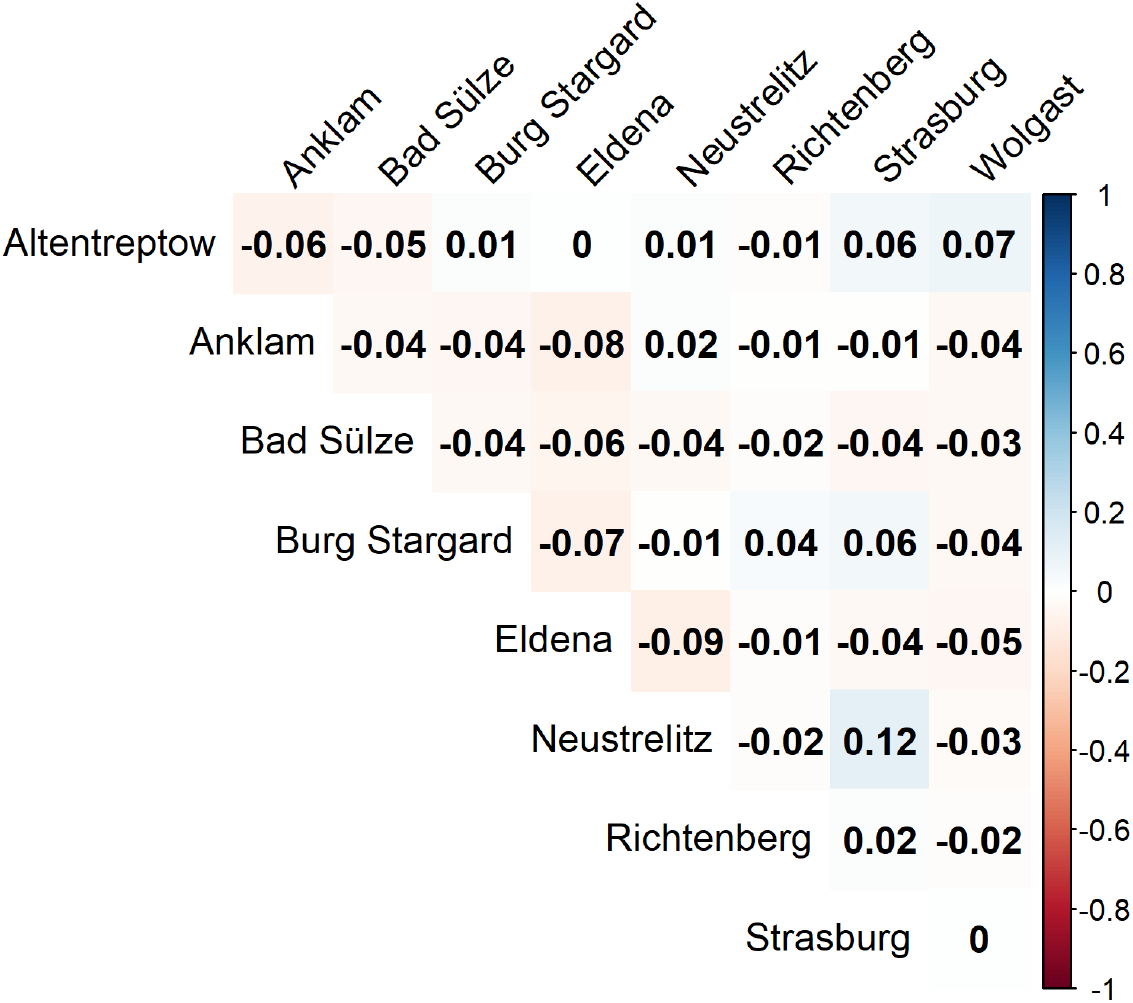
Correlation of MLG abundances across sampled hibernacula. Values and colour represent Pearson product-moment correlation indices. None of the correlations were significant (alpha = 0.05) following sequential Bonferroni-Holt correction of p-values.

### 3. Additional analyses on *P. destructans* MLG distribution between bats and walls in Eldena

#### 3.1 Minimum number of spores transferred from walls to bats (bottleneck size)

The determination of bottleneck size for different winters was obtained by simulating the transmission of spores from walls to bats using the available information on genotypic richness on bats and walls.

Specifically, the ‘Pool 0’ was composed of all SSIs isolated from the wall swabs of the Eldena hibernaculum (across all years, with MLG abundances exactly as observed from wall swabs in Eldena). We sampled (with replacement) different quantities of spores (*N* in the range 2-3100) from the ‘pool 0’ and obtained a new ‘Pool 1’. We then sampled 17.5% of spores from ‘Pool 1’ to take into consideration the germination rate calculated for European isolates (4 isolates; mean= 0.175; based on data in Fischer et al., 2020). For each bottleneck size, we then quantified the number of MLGs after the two successive samplings (i.e., for ‘Pool 2’; Figure S4). This procedure was repeated 1000 times for each bottleneck size, resulting in 1000 values of MLG richness for each value of *N* (see Figure S5).

**Figure S4.**
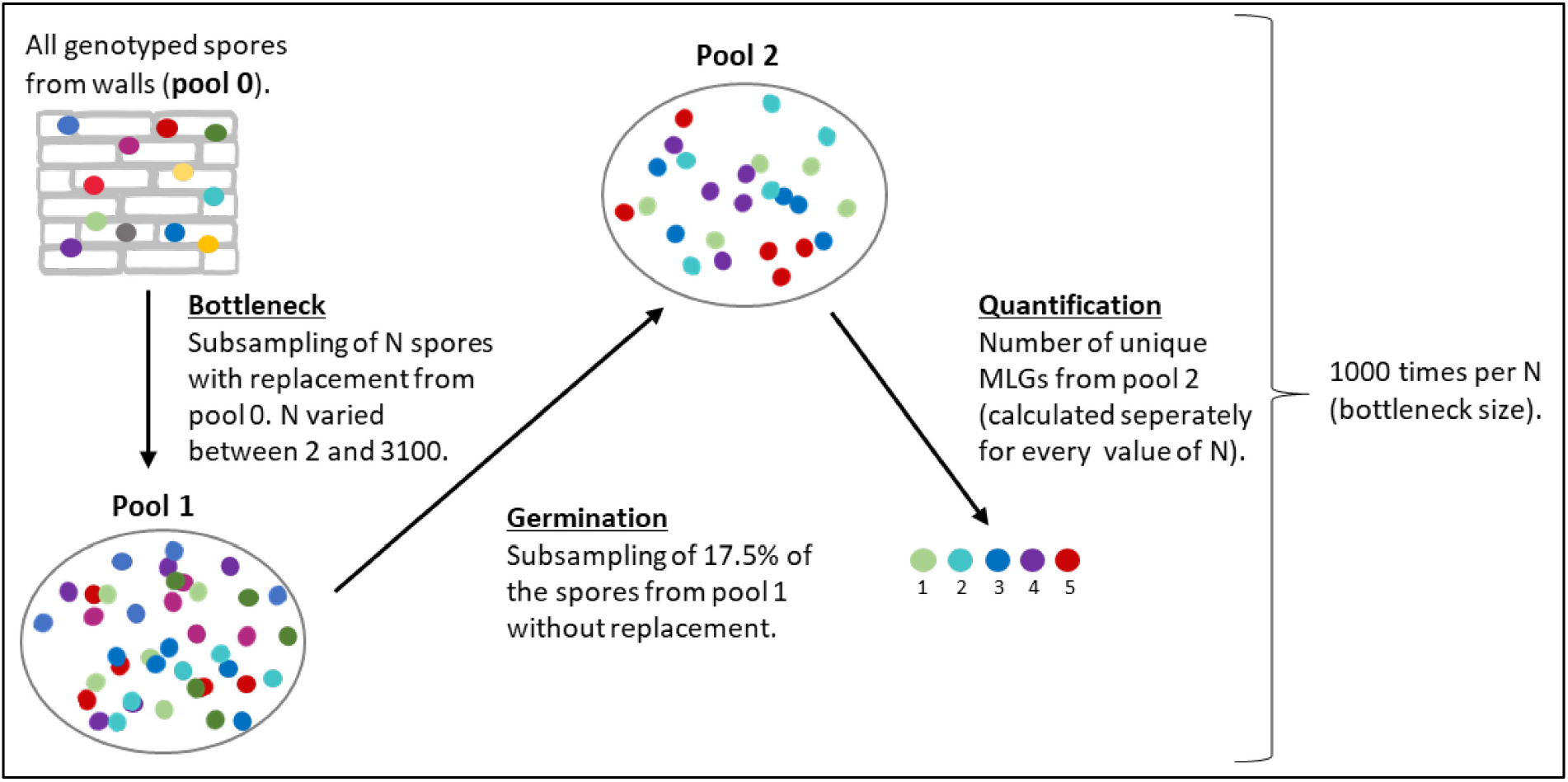
Schematic visualization of the simulation steps used to estimate the number of spores transferred from walls to bats early in the winter season. Coloured circles represent different MLGs.

**Figure S5.**
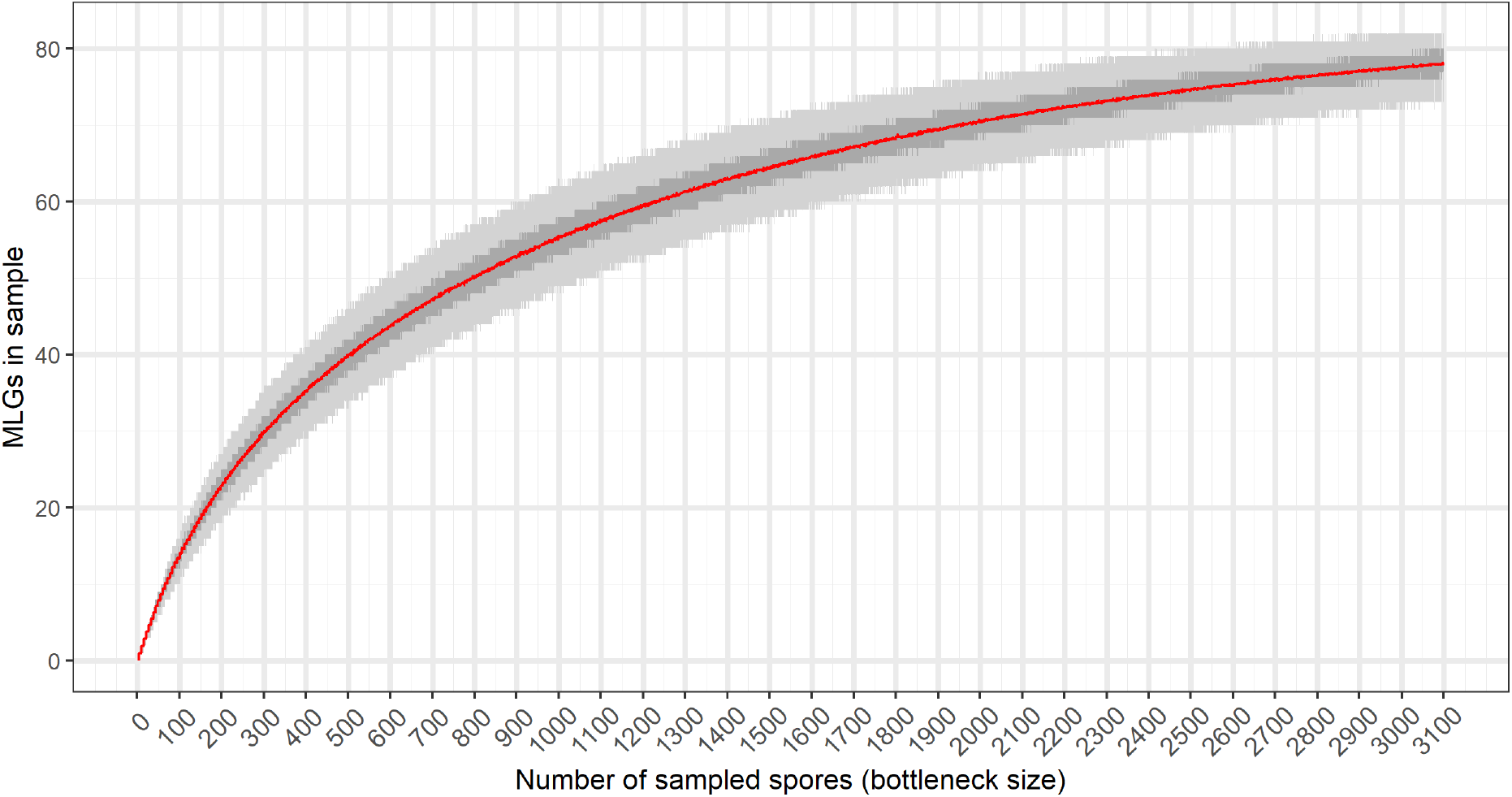
Number of unique MLGs (y-axis) for a given number of sampled SSI (x-axis; size of bottleneck, steps of 1 between 2 and 3100) from all SSI collected from walls. The red line shows the mean number of MLGs for each bottleneck size across 1000 runs, while the area shaded in dark grey and light grey show the 50% and 95% confidence interval, respectively.

We then compared the quantity of MLGs from all simulated bottleneck sizes with the genotypic richness (number of MLGs) isolated from bats in Eldena for different winters. To get more precise results we also used a predicted number of MLGs for bats in addition to the observed number (obtained via a Bayesian estimator classically used to estimate population sizes based on a single sampling session; Petit & Valiere, 2006; Puechmaille & Petit, 2007). This predicted number of MLGs accounts for MLGs that were present in the samples but had not been observed (e.g., not cultured and/or not genotyped). The bottleneck size (*N*) that gave the average MLG number closest to the observed/predicted number of MLG on bats was considered as the estimated number of spores transferred from walls to bats (to the entire sampled population of bats). To obtain an average number of spores passed from the wall reservoir to each single bat (Table S2), we then simply divided the determined matching bottleneck size (i.e., number of spores transferred to the entire sampled population of bats) by the number of body parts that were sampled (i.e., the number of samples) and multiplied by six, the number of body parts that are commonly infected by *P. destructans* (i.e., left/right ear, left/right wing, nose and uropatagium).

**Table S2:** Number of sampled body parts (i.e., samples; “N_Sample”_) per winter season for bat samples only. The numbers of spores transferred from walls to bats (overall; “Spores-all) as well as to a single bat (“Spores-bat”) were determined based on observed and predicted numbers of MLGs (“MLG”) and their match with genotypic richness observed for different bottleneck sizes.

| Winter season | $N_{\text{Sample}}$ | Observed | | | Predicted | | |
| --- | --- | --- | --- | --- | --- | --- | --- |
|  |  | MLG | Spores- all | Spores- bat | MLG | Spores- all | Spores- bat |
| Winter 14/15 | 139 | 74 | 2403 | 104 | 74 - 77 | 2324 - 2994 | 100 - 129 |
| Winter 15/16 | 31 | 34 | 375 | 73 | 35 - 48 | 381 - 741 | 74 - 143 |
| Winter 16/17 | 10 | 16 | 120 | 72 | 19 - 52 | 146 - 884 | 88 - 530 |
| Winter 17/18 | 53 | 37 | 435 | 49 | 37 - 41 | 422 - 540 | 48 - 61 |
| Winter 18/19 | 53 | 47 | 695 | 79 | 47 - 56 | 677 - 1065 | 77 - 121 |

#### 3.2 Testing for differences in genotypic richness using permutation test

To test for differences between genotypic richness on bats and walls in Eldena, we used a permutation test based on the numbers of unique MLGs.

For this we subsampled all SSI obtained from bat samples (N = 788) down to the sample size (total number of SSI) obtained from walls (N = 274). We then compared the number of unique MLGs in the subsample from bat swabs to the observed number of unique MLGs from wall swabs (thus having the same sample size).

The range of unique MLGs from 1000 subsamples of bat SSIs was 63 – 90 (Figure S6) with a mean of 76.5 and a median of 77. The observed number of unique MLGs from wall SSIs was 85 (at the same sample size). The number of unique MLGs from subset bat SSIs was therefore significantly lower than that observed from wall SSIs (One-sided permutation test, *df* = 999, *p* < 0.001).

**Figure S6.**
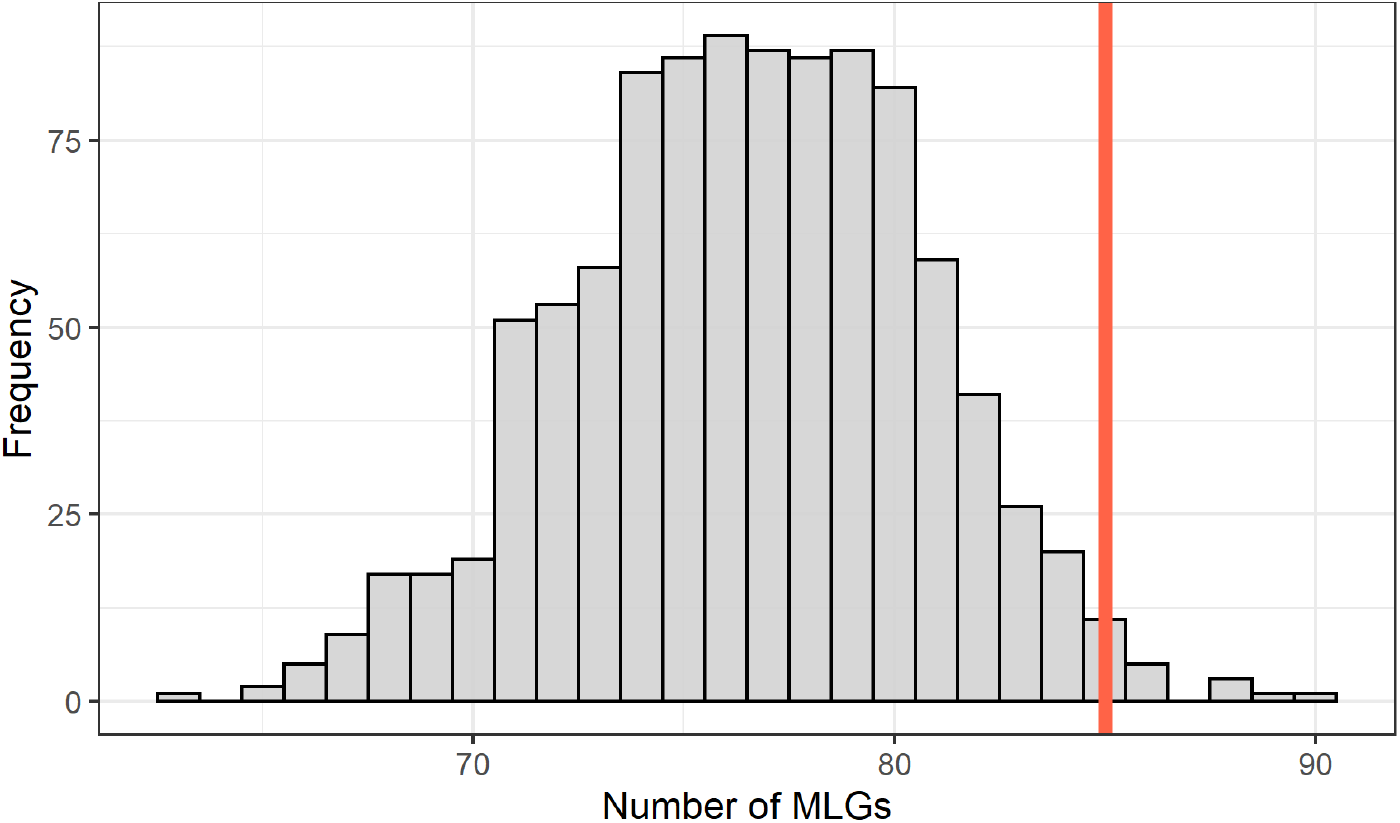
Histogram of number of unique MLGs in subsamples from bat SSIs (grey bars); sample size = 274; 1000 subsamples. Red line at 85 indicates the number of observed unique MLGs in wall SSIs (also with a sample size of 274).

### 4. Genotypic richness of *P. destructans* on bats and walls with 3 SSI per swab (Eldena)

We decided to show the results obtained from using 3 SSI per bat swab and 5 SSI per wall swab for all analyses on patterns of *P. destructans* between bat and wall in Eldena. This was done to obtain large sample sizes to best estimate genotypic richness overall as well as at each sampling event.

In the following, we present the results for analyses on genotypic richness between bats and walls under the same sample size (considering exactly 3 SSI per swab independent of the sampled substrate).

After excluding all samples for which we had obtained less than 3 SSIs, all swabs collected from bats yielded exactly 3 SSIs per swab. However, usually 5 SSIs were obtained from wall swabs resulting in the need to subsample the SSI from these swabs down to also obtain 3 SSIs. Because the selection of SSIs per swab could influence the observed genotypic richness, the subsampling of SSIs from wall swabs was done 1000 times to obtain estimates independent of which SSI combination was picked. The resulting dataset contained 150 SSI from wall samples and 699 SSI from bat samples.

#### 4.1 eMLG of SSIs from walls and bats (overall and temporal)

The eMLG for *P. destructans* SSI from bat swabs with exactly 3 SSIs per swab was 56.47 while the mean eMLG from wall swabs (subsampled to 3 SSIs per swab, 1000 runs) was 60.48 (at smallest shared sample size of 150; see Figure S7). Therefore, at the same number of SSIs per swab and independent of sample size per substrate, *P. destructans* SSI from walls have greater genotypic richness than those from bats.

In most cases, the mean eMLG of *P. destructans* from wall swabs was higher than the eMLG for SSI from bat swabs collected the same winter season (Figure S8). The exception to this was the winter of 2014/15, when genotypic richness on bats was slightly higher than that on walls.

**Figure S7.**
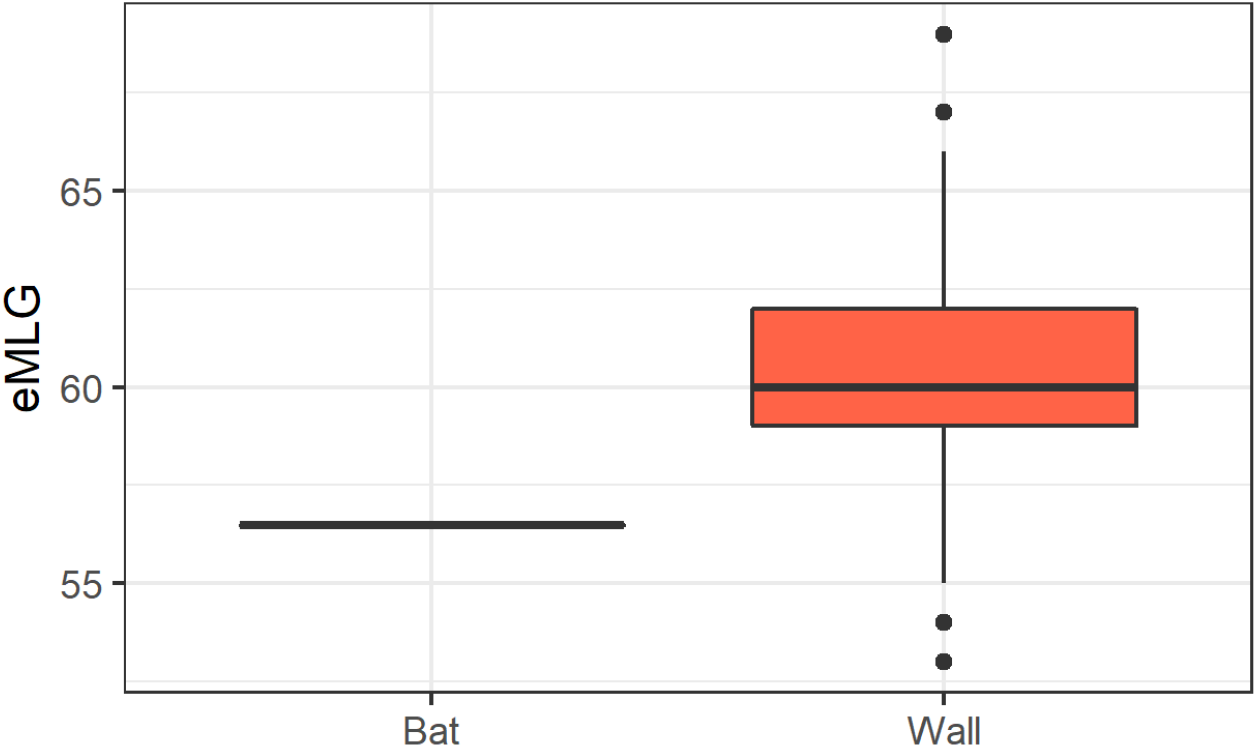
Genotypic richness of *P. destructans* from bat and wall samples at exactly 3 SSIs per swab. The smallest shared sample size was 150 (SSIs from wall swabs). Bars show interquartile range between the 25^th^ and 75^th^ percentile; line shows the median; whiskers represent the largest and smallest value within 1.5 times the interquartile range; points show outliers between 1.5 and 3 times the interquartile range.

**Figure S8.**
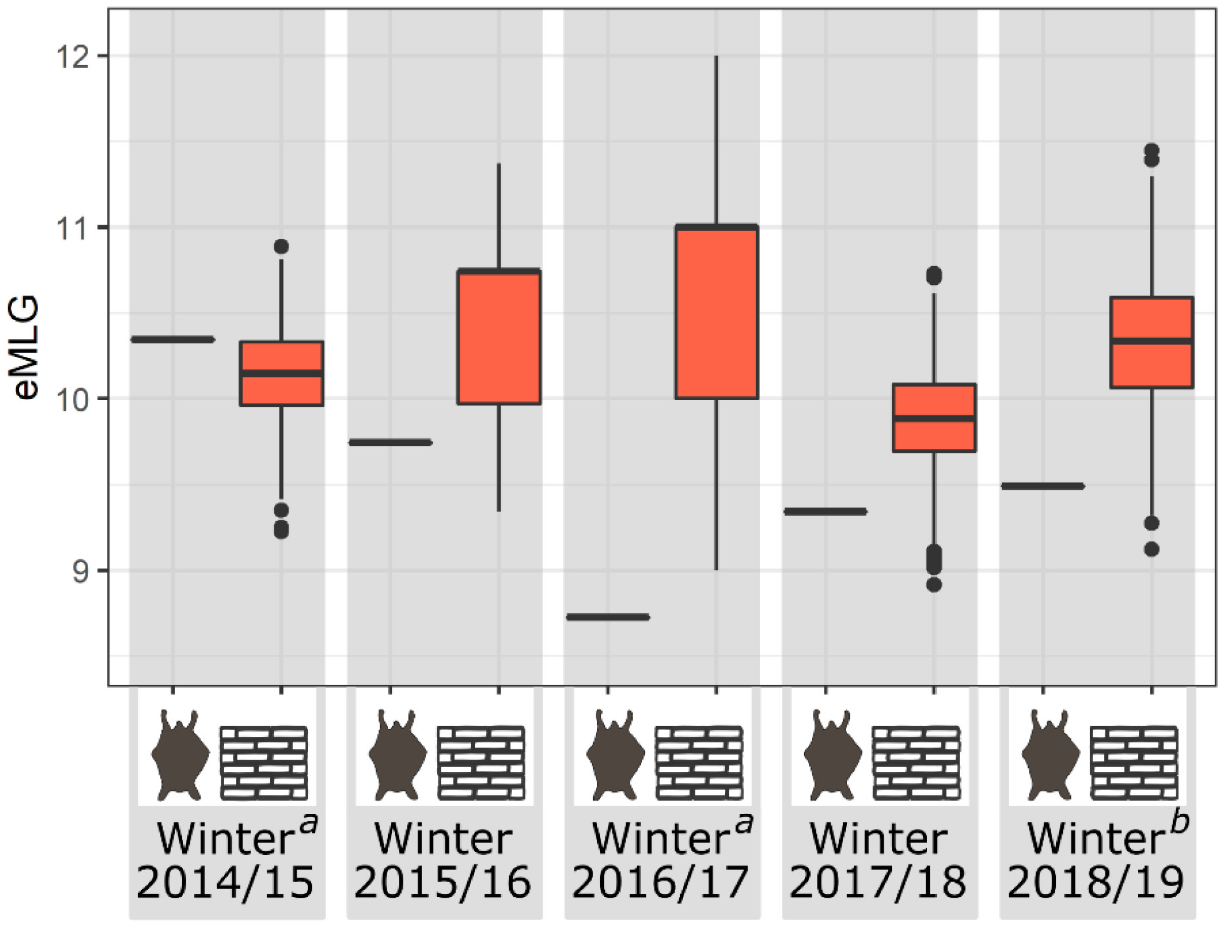
Genotypic richness (eMLG) of *P. destructans* for each substrate (indicated by bat and wall symbols) by winter season at a smallest shared sample size of 12 (wall winter 16/17). Bars show interquartile range between the 25^th^ and 75^th^ percentile; line shows the median; whiskers represent the largest and smallest value within 1.5 times the interquartile range; points show outliers between *^a^* contains isolates from wall sampling in April only. *^b^* contains isolates from wall sampling in October only.

#### 4.2 Testing for differences in genotypic richness using permutation test

This analysis follows the same principle as shown in section S3.2 but on the dataset of wall and bat SSI subsampled to exactly 3 SSI per swab. To quantify the genotypic richness on bats and walls in a permutation test (in addition to the eMLGs obtained in S4.1), we subsampled the SSI obtained from bat samples (N = 699, at exactly 3 SSI per swab) down to the sample size obtained from wall samples (N = 150, at exactly 3 SSI per swab). We then compared the obtained number of unique MLGs in the subsample to the observed number of unique MLGs from wall swabs (i.e., at the same sample size and the same number of SSI per swab). This resulted in 1000 values for bat swabs (which did not have to be previously subset to obtain 3 SSI per swab) and 1.000.000 values for wall swabs (which were first subset to 3 SSI per swab in 1000 runs before running this permutation test).

The mean number of unique MLGs from bats was 56.47 while the mean number of unique MLGs from wall swabs was 60.50 (Figure S9). At the same sample size, samples from the walls of hibernacula show a greater number of unique MLGs than samples collected from bats (Two-sample Kolmogorov-Smirnov test, *D* = 0.48, *p* < 0.001).

**Figure S9.**
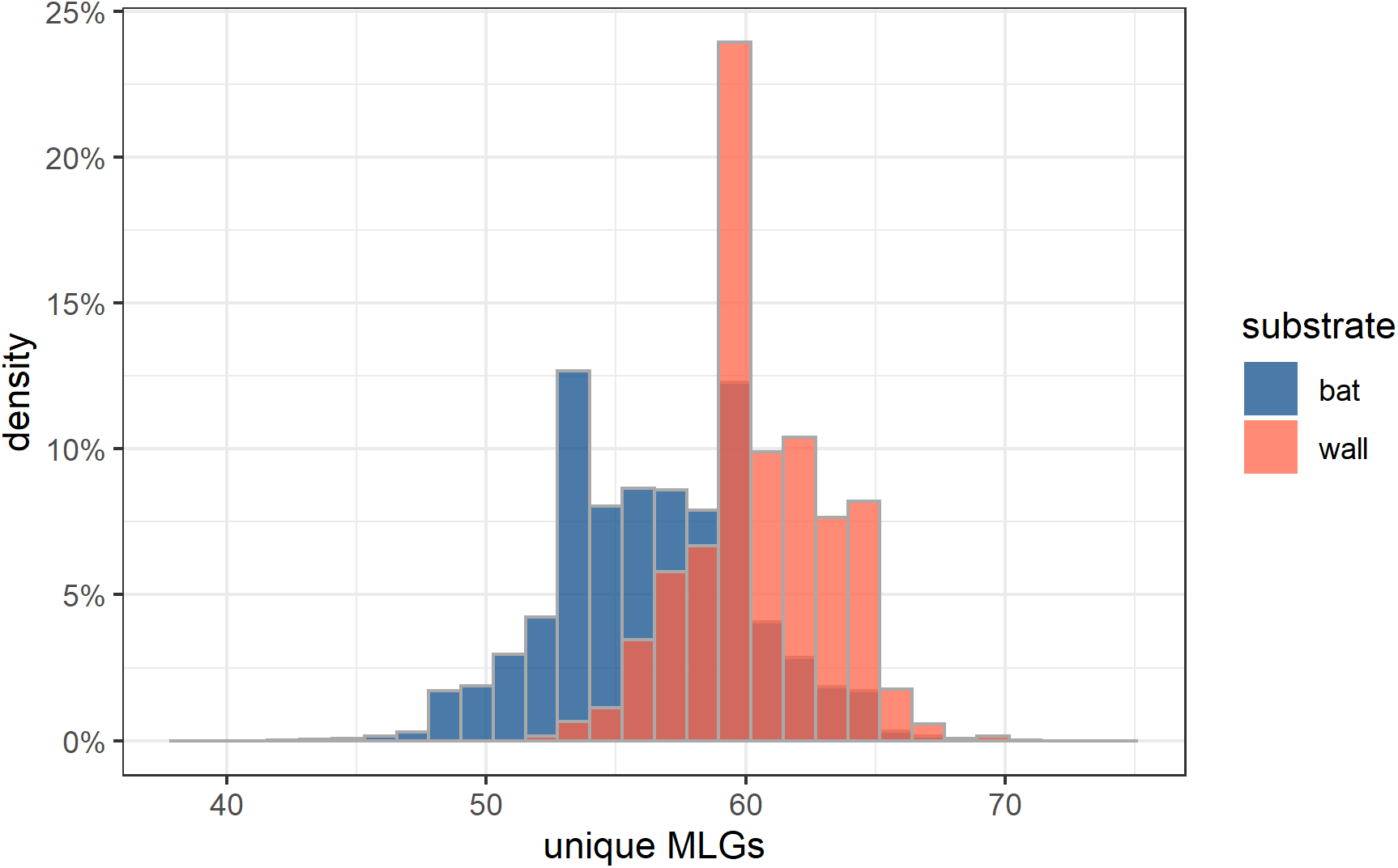
Density distribution of the number of unique MLGs following subsampling of bat SSI (N=699) to the number of SSI from walls (N = 150) with exactly 3 SSI per swab sample.

